# A simple CPG-based model to generate human hip moment pattern in walking by generating stiffness and equilibrium point trajectories

**DOI:** 10.1101/737031

**Authors:** Alireza Bahramian, Farzad Towhidkhah, Sajad Jafari

## Abstract

Equilibrium point hypothesis (its developed version named as referent control theory) presents a theory about how the central nerves system (CNS) generates human movements. On the other hand, it has been shown that nerves circuits known as central pattern generators (CPG) likely produce motor commands to the muscles in rhythmic motions. In the present study, we designed a bio-inspired walking model, by coupling double pendulum to CPGs that produces equilibrium and stiffness trajectories as reciprocal and co-activation commands. As a basic model, it is has been shown that this model can regenerate pattern of a hip moment in the swing phase by high correlation (*ρ* = 0.970) with experimental data. Moreover, it has been reported that a global electromyography (EMG) minima occurs in the mid-swing phase when the hip is more flexed in comparison with the other leg. Our model showed that equilibrium and actual hip angle trajectories match each other in mid-swing, similar to the mentioned posture, that is consistent with previous findings. Such a model can be used in active exoskeletons and prosthesis to make proper active stiffness and torque.

## Introduction

Equilibrium point hypotheses (EPH) and its extension, referent control theory [1, 2], are effective theories in responding to the question of “how central nerve system (CNS) controls the body?” [3, 4]. These theories explain how the CNS can release from complex calculations to control movements. According to EPH, at the muscle level, the CNS only determines a specific threshold (*λ*) for muscle [5, 6]. If actual muscle length goes beyond the defined threshold, EMG activity emerges as a result of the gap between actual and threshold muscle lengths. To control the joint, two commands are defined by the CNS: reciprocal (R) command and co-activation (C) command [7]. If joint is considered as rotational spring, R and C commands determine equilibrium point and stiffness of the joint, respectively. In accordance with the referent control theory, for multi-joint system (e.g. leg), it is assumed that initially, global R and C commands are set by CNS, then, by few-to-many mappings, r and c commands for various joints are defined. Finally, *λ*s As for agonist and antagonist muscles emerged from these r and c by another mapping [8]. This hierarchical system is effective for making coordination between different joints and muscles [9]. One of the powerful experimental evidence for this hierarchical referent system is the global EMG minima [10]. When the referent (R) and actual configurations approach meet each other, a minimum EMG activity occurs for most of the muscles involved in the task [10]. This phenomena has been confirmed by many studies on various tasks such as walking and jumping [10], monkeys head movement [11], Jete in ballet dancers [12], and hammering [13]. It is shown that during stepping forward, a global EMG minima occurred during mid-swing when the hip flexion of the swing leg is slightly more than the other leg [14].

On the other hand, the behavior of the joints as a rotational spring have commonly been used for investigating the relationship between torque(s) and angle(s). These studies are applicable to design anthropomorphic biped robots [15–17], exoskeletons [18, 19] and prosthesis [20, 21]. Moreover, the slope of the relationship between toque(s) and angle(s) has been defined as “quasi-stiffness” which is widely used to design these systems [22]. There are some research on estimating quasi-stiffness for the hip joint in the swing phase [23, 24].

Moreover, it has been believed that some neural circuits (so-called CPGs) located in the spinal cord [25, 26], generate motor commands for the limbs in human and animals locomotion [4, 27]. CPGs can produce rhythms of motion autonomously, however, they are affected by ascending brain commands and also sensory feedbacks [28]. Some models considered the CPGs as a generator of rhythmic motor commands [29, 30]. Some of CPG-based models represent hierarchical controller inspired by the biological structure of human motion [31], especially by coupling reflexes to the CPGs to control biped motion [32, 33]. One of the effects of reflexes on CPGs is known as phase resetting [34]. It refers to reset the rhythmic activity of CPGs produced by perturbations [35] or events such as foot-ground contact [36], which can be seen in human [37] and animals [38, 39]. In addition, several humanoid biped models showed the effectiveness of ground foot contact phase resetting on robustness and stabilization of model [36, 40, 41].

Biped rhythmic motion such as walking involves lots of muscles and different joints by complex morphology. However, some simple and conceptual models provide valuable insight for better understanding of these complex process [42, 43]. For example, global behavior of running could be explained by a simple spring model for the leg, with certain equilibrium and stiffness [44]. Also, nonlinear joint stiffness that contributes to make a global linear spring behavior of the leg during running, could be explained by modeling ankle and knee as rotational spring [45]. Therefore, such these simple models are useful to investigate dynamics governing such rhythmic biped motion [46, 47]. Double pendulum model was used in this study as a conceptual model which has been also frequently used to investigate human biped walking [48, 49].

The main aim of this study is to explain how the pattern of the hip moment in the swing phase can be expressed by a combination of CPGs and EPH. Therefore, we first introduced our model and then the experimental data applied to design CPGs and estimate model parameters. Finally, the model performance is evaluated by comparing with experimental data.

## Materials and method

### Biped model

Double pendulum model equations were used in this study for single and double support phase [50, 51]. Eq.(1) and Eq.(2) are used as a model of single support phase:

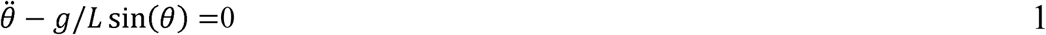

Eq.(3) is the condition for the end of the single support phase:

Impact of swing leg to the ground affects both leg velocity at the beginning of double support phase. Also, switch between swing and stance legs together with ankle push-off are two other important events of double support phase. These events are modeled in Eq.(4):

In Eq.(4), (-) and (+) note states values before and after double support phase. Also, represent the amplitude of the push-off impact. Schematic of single support and double support phase model is illustrated in Fig.1.

**Fig. 1.**
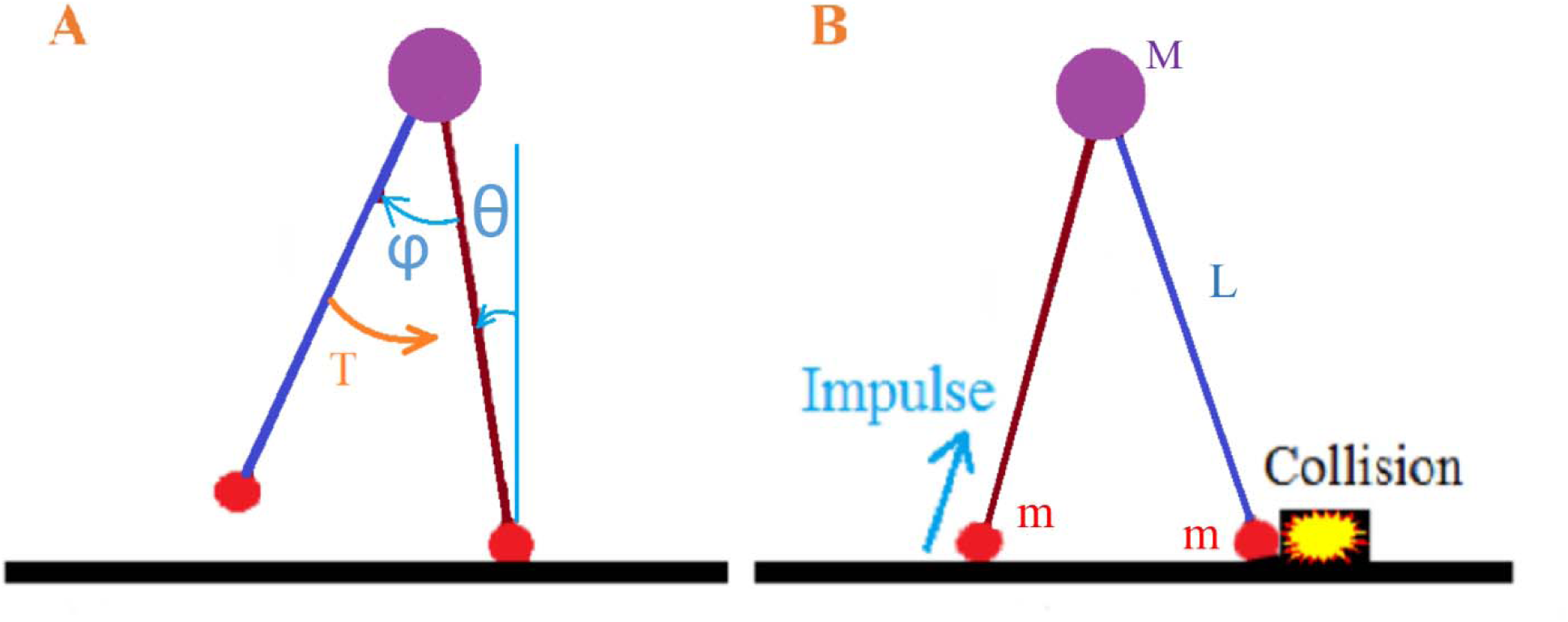
Representation of single and double support phase models. A) Single support phase; hip torque is applied to the swing leg from the assumptive torso. B) Double support phase; push-off *Impulse* is applied to the double pendulum. Moreover, ground collision and alternated switching between two legs were assumed in Eq.(5).

The values considered for model parameters are mentioned in table 1.

**Table 1.** considered value for model parameters.

| Parameter | Value |
| --- | --- |
| $M$ | 30 |
| $m$ | 5 |
| $L$ | 0.75 |
| $g$ | 9.8 |

### Experimental data

Young group walking data (9 males, 11 females; mean age: 10.8 ± 3.2 years; height: 1.47 ± 0.20 m; body mass: 41.4 ± 15.5 kg) was used in this study [52] (for more detail see [52]). Hip angle and hip torque data for normalized stride are plotted in Fig.2 and Fig.3, respectively.

**Fig. 2.**
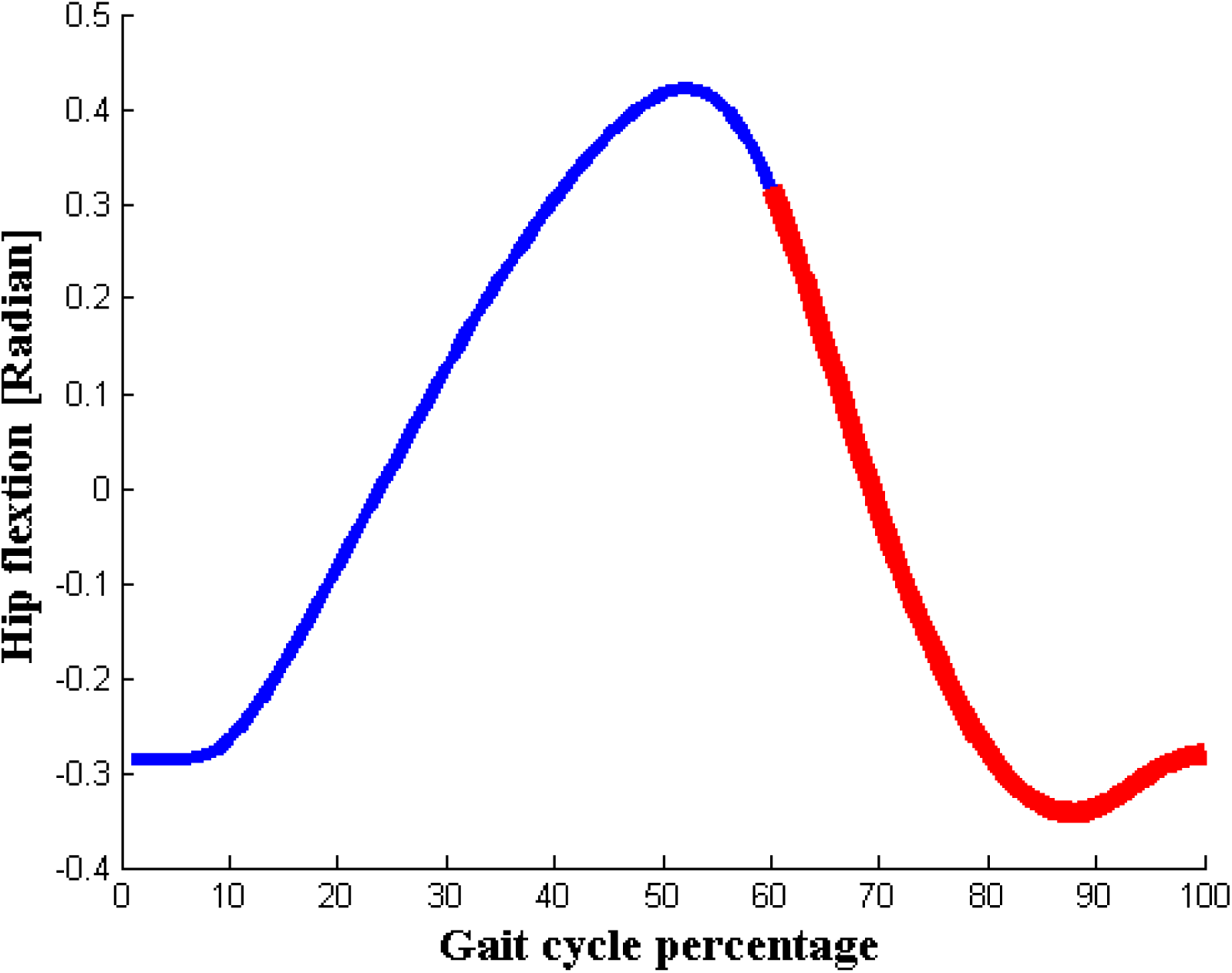
The angle of hip joint for normalized stride from heel strike to the next heel strike is plotted. Hip angle during the swing phase is shown by red line.

**Fig. 3.**
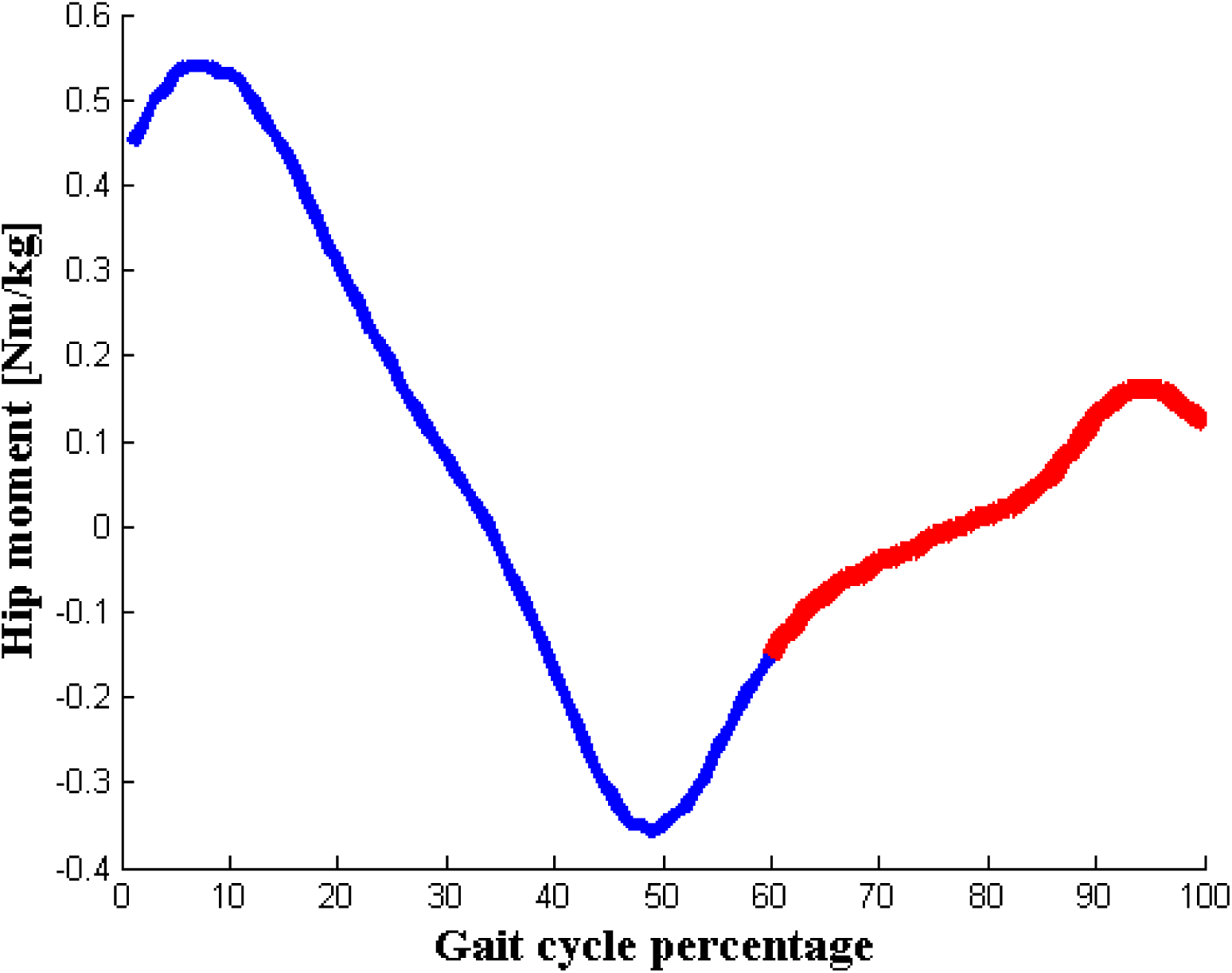
The hip moment for normalized stride from heel-strike to heel strike is shown. Duration of swing phase is drowned by red line.

### CPG design

Based on the equilibrium point hypothesis, the torque that applied to the swing hip is defined as Eq. (5).

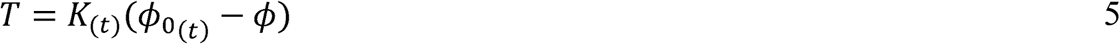

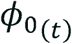 is equilibrium and *K*_*(t)*_ is stiffness of hip joint trajectory as a function of time. Pattern of 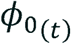 and *K*_*(t)*_ are considered to generate by CPGs. feasible patterns for 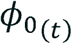 and *K*_*(t)*_ is estimated by the pattern of the hip moment during the swing phase, for the design of CPGs (Fig.4). In the early swing phase, the torque of hip tries to force the leg forward, but after about mid-swing, it pushes the leg backward. So, at the beginning of motion, *ϕ* must be backward in comparison with its equilibrium trajectory 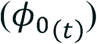. After mid-swing, dynamics of motion and velocity of swing leg throws *ϕ* ahead related to 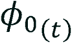. The increase of *K*_(*t*)_ leads to an increase in the hip moment. Therefore, high or low values of moment could be a consequence of the high or low value of stiffness (besides the difference between the angle and its equilibrium trajectory).

Pattern of and are designed in order to generate by simple CPGs using Eq.(6) and Eq.(7).

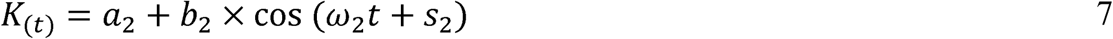

*a*_*i*_, *b*_*i*_, *ω*_*i*_, and *s*_*i*_ are setting parameters of CPGs. These CPGs patterns reset (t →0) in the double support phase, when the mentioned condition in Eq.(3) is established. Consequently, in this way, CPGs are coupled with the biped model. First, the parameters of CPGs are configured manually and then they are optimized by the genetic algorithm to achieve the most correlation between experimental hip moment and *T* of the model.

**Fig. 4.**
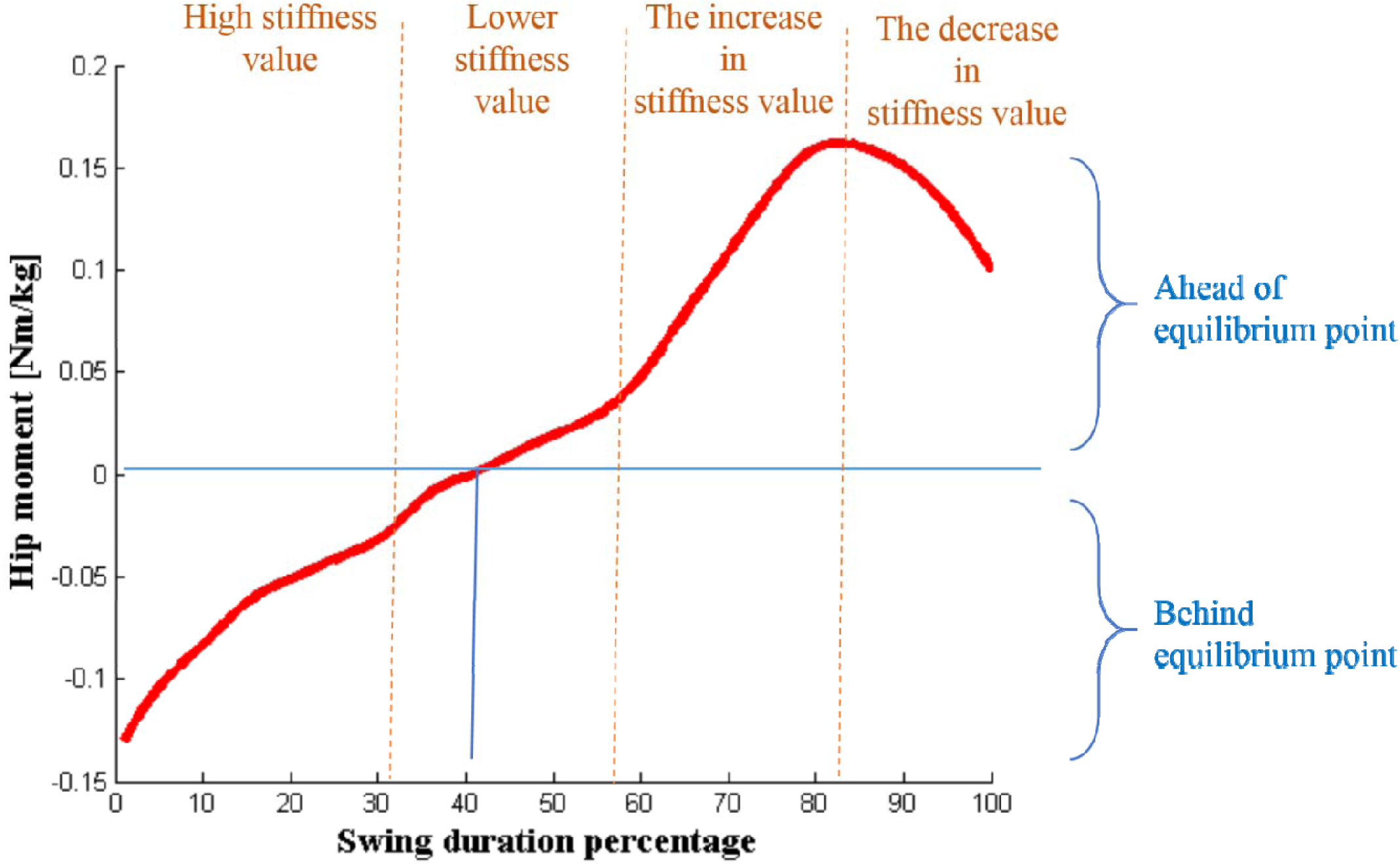
The normalized hip moment during swing phase. Sign of moment value determines when joint is behind or ahead of its equilibrium trajectory. The high value of moment also can be as a result of the high value of assumed stiffness. In addition, the decrease of the moment at the end of motion can be caused by the decrease of stiffness value.

## Result

By setting optimized parameters of the CPGs, after the transition time, the dynamic behavior of *θ* and *ϕ* are displayed in Fig.5. Stability of the model is confirmed by perturbing model and using Poincare section method [50, 53, 54]. The hip angle of swing leg of model correlated strongly with experimental data (*ρ*= 0.9220).

**Fig. 5.**
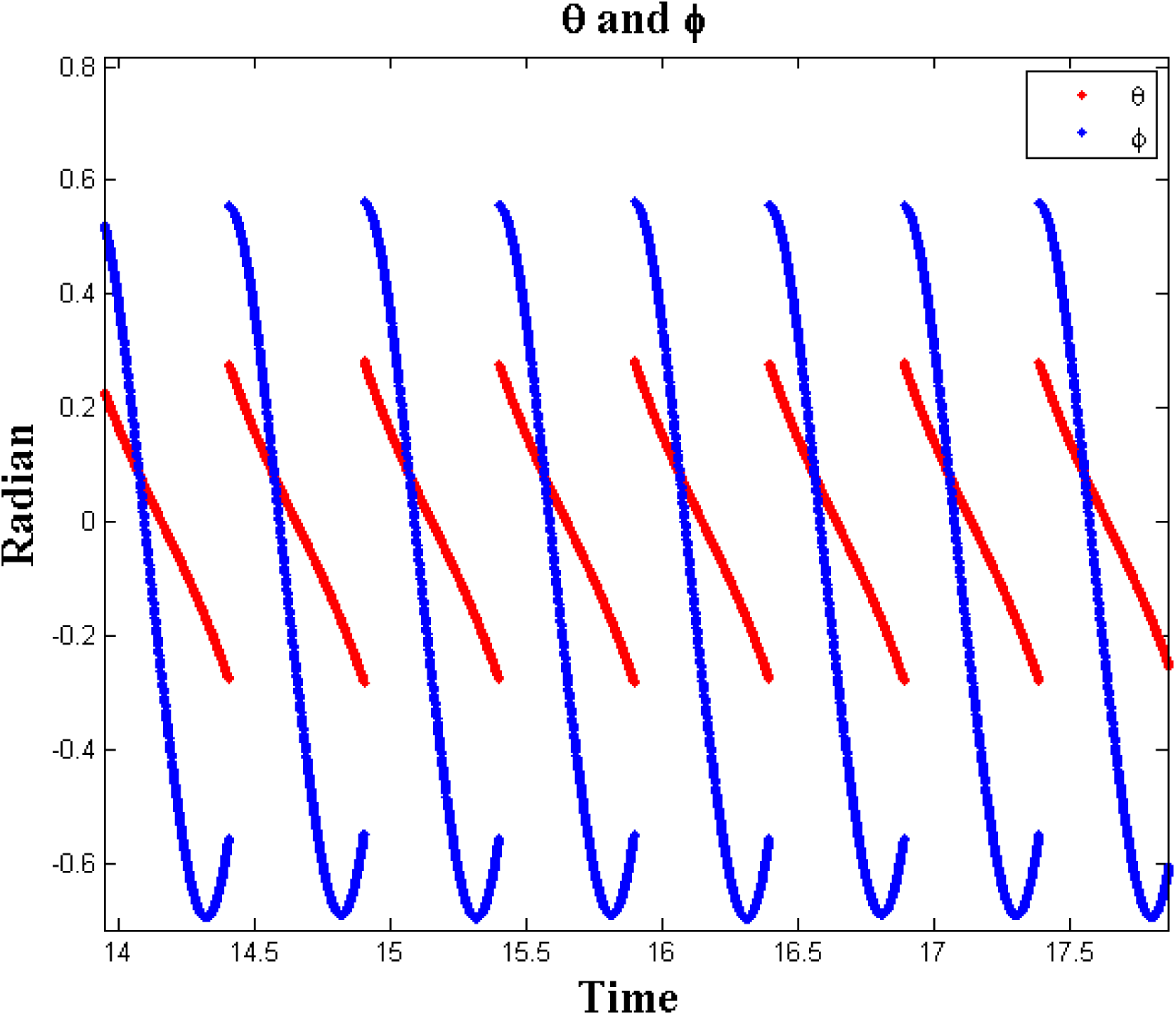
An illustrative behavior of the double pendulum model. *θ* and *ϕ* are drowned by red and blue line, respectively.

Final pattern of 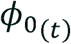 and *K*_(*t*)_ are demonstrated in Fig.6 and Fig.7. As can be seen, 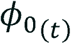 changes monotonically in the forward direction. Also, the pattern of *K*_(*t*)_ shows a decrease of stiffness value in the mid-swing and at the end of the swing phase.

**Fig. 6.**
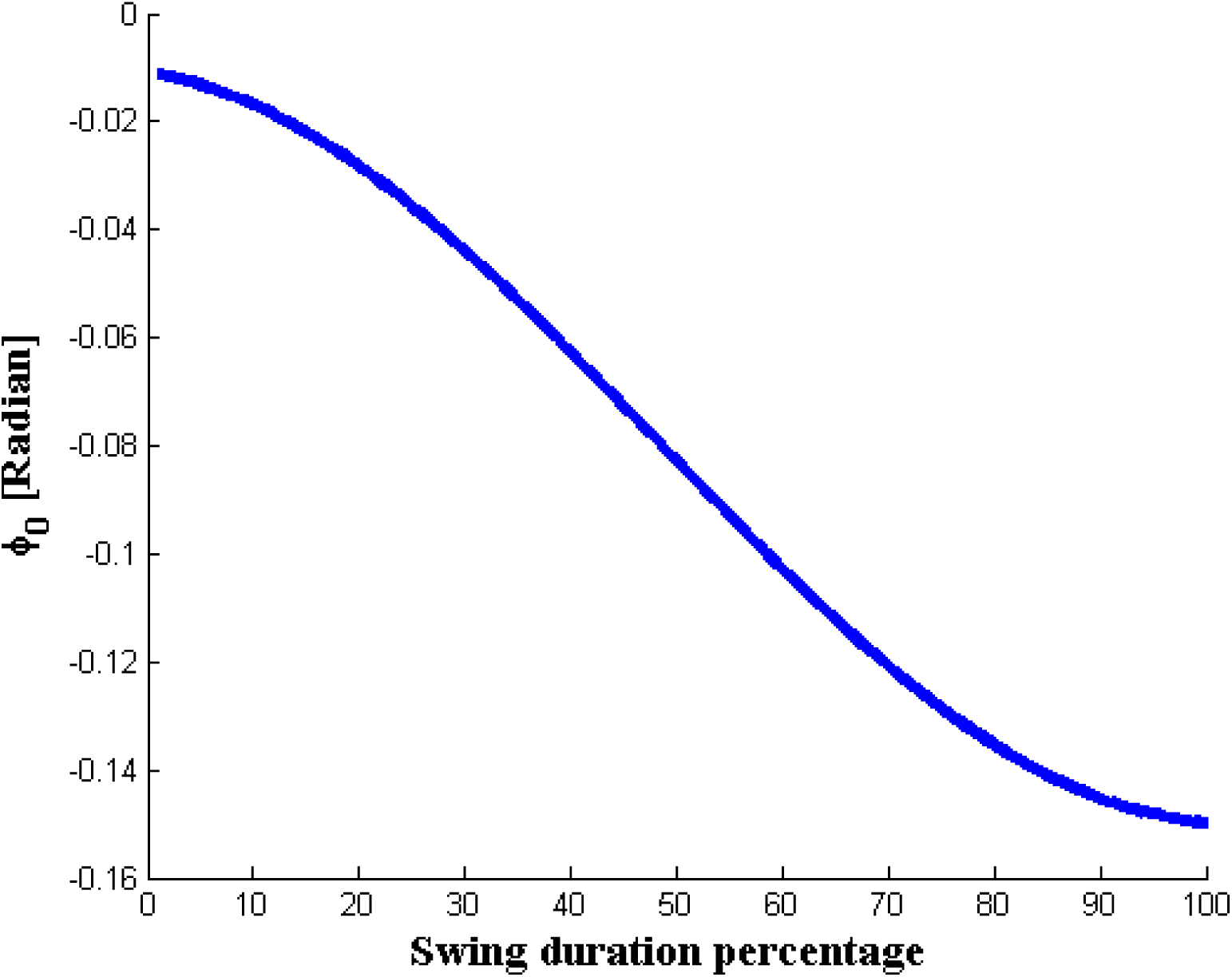
Normalized 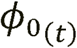 during the swing phase of the model at the steady-state.

**Fig. 7.**
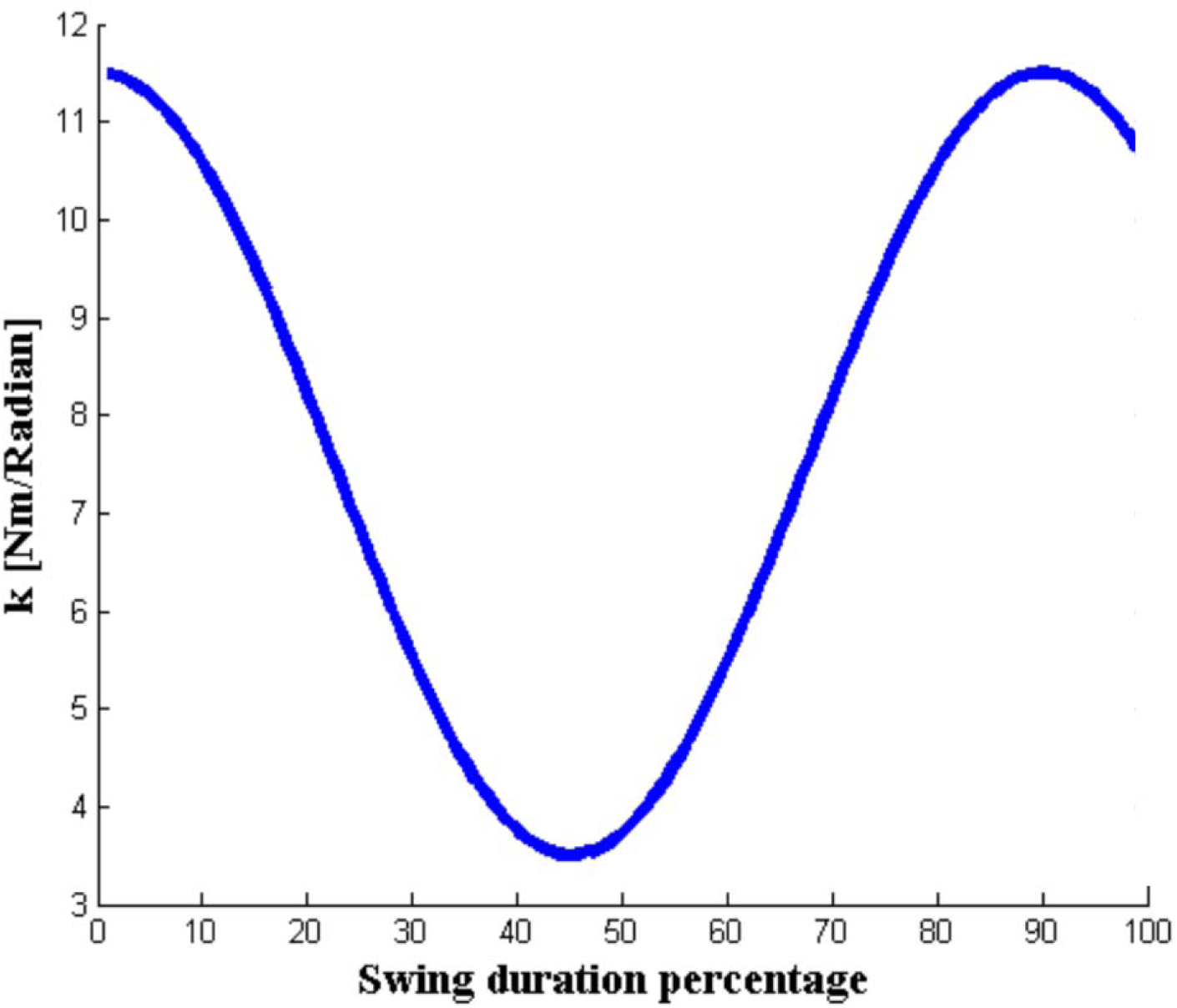
Normalized during the swing phase of the model at the steady-state.

pattern during the swing phase is shown in Fig.8. There was a high correlation () between the model and experimental data for the hip moment. is equal to zero when is equal to.

**Fig. 8.**
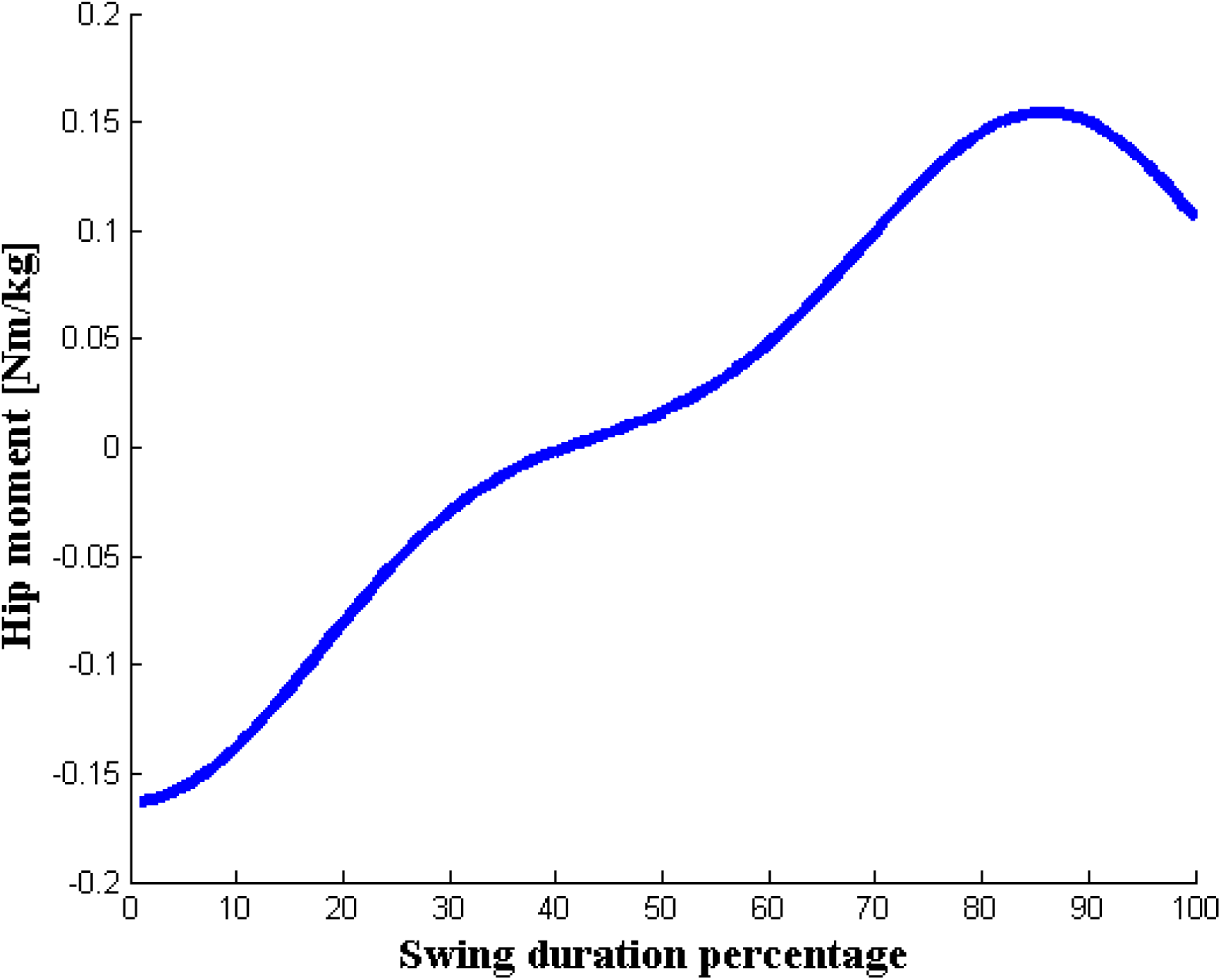
Normalized *T* during the swing phase of the model at the steady-state. It has high correlation *ρ* = 0.9790 with experimental hip moment data.

## Discussion

In the present study, we used the equilibrium point hypothesis, as a well-known theory of motor control to investigate the hip moment during the swing phase. Double pendulum model was applied to the model of walking. We used simple CPGs to generate equilibrium and stiffness trajectories and coupled them with the double pendulum model. The bio-inspired model has been introduced to generate hip moment pattern. Our result showed that there was a high correlation (*ρ* = 0.9780) between the model and experimental data for the hip moment of the swing leg. Such conceptual model helps to get a better vision about complex dynamical motion without confusing by involving redundant details.

Some types of CPGs are introduced that designed to produce appropriate torques for joints [55, 56]. Answering to the question of “what is sufficient and best language to clarify control in the neuromuscular system”, force control theory has faced many serious criticisms [57, 58]. Instead, referent control of movement has been suggested to explain how the nervous system controls movements [59]. There are few research assumes the CPGs as generators of equilibriums and stiffnesses trajectories [60, 61]. These kinds of CPGs which were also used in this study are more suitable as a bio-neuromuscular model.

After optimization to achieve the most correlation between the hip moment of experimental data, a simple monotonic pattern for equilibrium trajectory 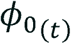 was revealed. The monotonic and non-monotonic pattern for equilibrium trajectory shift are assumed for the different motor task [62, 63]. Our model revealed that only simple monotonic pattern for equilibrium trajectory of the hip is sufficient to generate hip moment pattern during swing phase.

As mentioned in the literature, a global EMG minimum occurs during stepping forward, in the mid-swing phase, when swing leg hip that passes the other leg in the stance phase, stays on a little ahead [14]. The proposed model showed the same results.

Several studies tried to extract the relationship between torques and angles of the hip [23, 24] or other lower body joints [64, 65] by assuming linear slop as quasi-stiffness between moments and angles. These researches are useful to design passive exoskeleton or prosthesis [66, 67]. However, active exoskeletons or prosthesis are capable to use a more advanced approach to generate required torques [68, 69]. These conceptual models are suggested to use as the origin of active stiffness generation. Moreover, their ability to model movement dynamics for generating an appropriate pattern for joints moment beside their fewer calculation demands in comparison with complex multi-link models makes them a proper option to this function.

## Conclusion

In this work, a bio-inspired model is designed for walking that employs concepts of CPG and referent control theory. In this model, R and C commands were built by CPGs. Hip moment pattern emerged from coupling CPGs to the model with high correlation (*ρ* = 0.9780), as compared to experimental data. In addition, according to our results, the simple monotonic motion of equilibrium trajectory can be sufficient for hip in the swing phase of walking. In this model, equilibrium trajectory and the angle of the hip have similar values in the mid-swing, when hip of the swing leg was ahead of the other leg in the stance phase. It is interesting to note that, in this posture, a global EMG minima is reported in the literature that confirms the result of our model experimentally. This kind of model is useful for emulating biped walking in some field of biomedical engineering.

